# Non-coding DNA dictates cell size and impairs fitness by sequestering RNA polymerase

**DOI:** 10.64898/2026.07.10.737727

**Authors:** Marianna E. Estrada, Shin Ohsawa, Gloria Sancho-Andrés, Ning Lu, Weihong Qi, Silvio Silva Lima, Jan Skotheim, Gabriel E. Neurohr

**Affiliations:** Institute of Biochemistry, Department of Biology, ETH Zürich, Zürich, Switzerland; Functional Genomics Center Zürich, Zürich, Switzerland; Department of Biology, Stanford University, Stanford, USA; Chan Zuckerberg Biohub, San Francisco, USA

## Abstract

Genome size varies greatly between organisms, largely because of varying amounts of non-coding DNA. In multicellular organisms, non-coding DNA can make up 99% of the genome, while unicellular organisms have little non-coding information. How non-coding DNA affects cell physiology remains a key question in understanding genome evolution. Here, we developed a system to conditionally accumulate large amounts of non-coding DNA in budding yeast cells. We show that cells adjust their size to DNA content, regardless of its coding capacity. The direct coupling of cell size to DNA content occurs independently of the known cell size control and DNA damage checkpoints and explains the long-observed correlation between cell size, genome size, and ploidy. Furthermore, we find that excess non-functional DNA compromises cell fitness and renders them hypersensitive to transcription inhibitors. Providing a potential explanation for this observation, we uncover that the transcription machinery binds non-coding DNA and is titrated away from coding genes, resulting in a reduced overall concentration of yeast transcripts. We propose that the inability of yeast to repress non-coding DNA increases the fitness cost associated with non-functional DNA and prevents its expansion in the yeast genome on evolutionary timescales.

## Main

Genome size, structure, and organization vary greatly between organisms. This variability results in part from different genomes having different coding capacities. Human genomes, for example, contain about four times as many protein-coding genes as the genome of the unicellular eukaryote *Saccharomyces cerevisiae* ^1,2^. This number appears small considering that the human genome is 250 times larger than the yeast genome ^1,2^. Variability in genome size therefore originates mostly from the presence of non-coding DNA ^3,4^, which can vary even between closely related species ^5^. In multicellular organisms, non-coding DNA can make up 99% of the genome ^6,7^, indicating that, in these organisms, it is well tolerated and it serves important cellular functions. Indeed, DNA regions that are not coding for proteins have been shown to modulate gene expression and genome homeostasis ^8^.

In contrast to the large and variable genomes of multicellular organisms, unicellular organisms tend to have small genomes. In Archaea and Bacteria, about 90% of the genome is protein coding ^9^. In *S. cerevisiae*, 72% of the genome codes for proteins, and the genome contains few introns, short intergenic regions, and little non-coding DNA ^2,7^. This gene-dense structure suggests that in unicellular eukaryotes, there is a strong selective pressure favoring the loss of non-coding DNA.

The physiological consequences of gene copy-number alterations have been broadly investigated, particularly in the context of whole-genome duplications (polyploidy) and chromosome copy-number aberrations (aneuploidy) ^10–12^. In contrast, how non-coding DNA affects organismal physiology remains a key open question in understanding genome evolution.

Interestingly, genome size variation is tightly correlated with changes in cell size ^13^. This correlation can be observed between cell types of the same species containing different genome copy numbers (ploidy) ^14^. Importantly, cell size and DNA content are also coupled when comparing different species with variable genome sizes ^15–17^ and even if the variation arises mostly from non-coding DNA ^18^. Several hypotheses might address why cell- and genome size co-evolve ^19^. Replication of expanded genomes might require larger cells to provide sufficient building blocks for DNA and chromatin synthesis. Alternatively, cell size could be under direct selection, and the expansion or shrinking of non-coding DNA might represent an evolutionary adaptation to altered size. The causality, mechanistic basis, and functional relevance of the coupling between DNA content and cell size remain enigmatic.

To investigate how non-coding DNA affects cell physiology, we developed an experimental strategy to accumulate up to one genome equivalent of human DNA in budding yeast cells. We find that this expansion of non-functional DNA leads to a proportional increase in cell size. This coupling does not rely on the established cell size control networks, demonstrating that the coupling of cell size and DNA content is direct and independent of gene dosage. In addition, non-coding DNA impairs cell fitness by titrating the transcription machinery away from endogenous genes. Our results therefore provide a mechanistic explanation for why non-coding DNA has been eliminated from the genomes of fast-growing unicellular organisms.

## Results

### A conditional system to accumulate large amounts of non-coding DNA in budding yeast

To determine the physiological consequences of harboring non-coding DNA in the genome, we set out to experimentally increase the amount of non-coding DNA in budding yeast cells (**Fig. 1a**). To this end, we conditionally inactivated the centromere of an 850 kb large yeast artificial chromosome (YAC-L) harboring human DNA with a similar sequence composition to yeast DNA (**Fig. S1a**). Centromeres were inactivated by placing the strong inducible *GAL1* promoter in front of them. Promoter activation by the addition of galactose leads to conditional centromere inactivation and retention of both sister chromatids in the mother cell during mitosis ^20,21^ (**Fig. 1b-d**).

**Figure 1.**
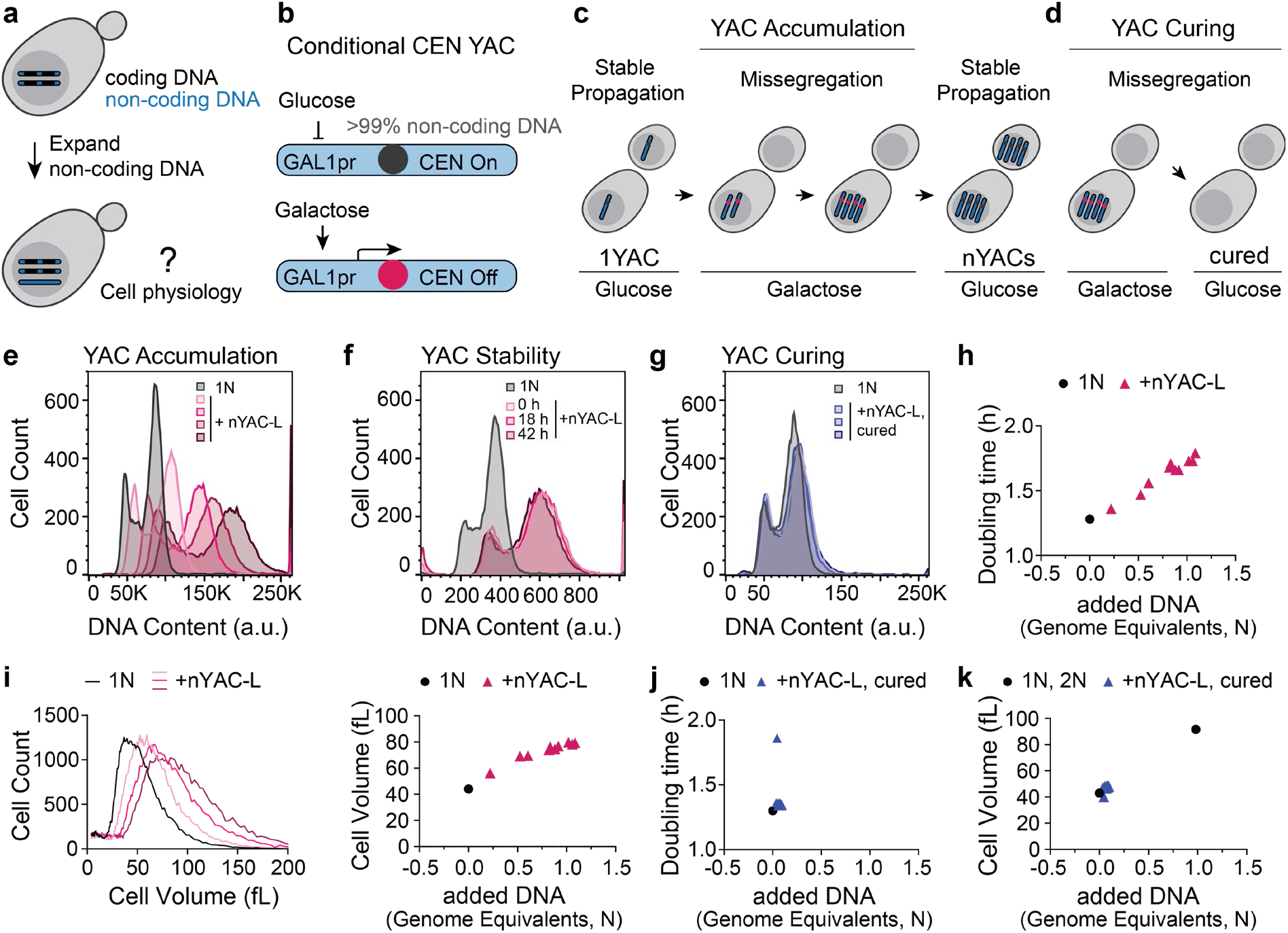
Non-coding DNA accumulation influences cell size and impairs cell fitness. **a**, Schematic illustrating the aim of this study to investigate how expansion of non-coding DNA affects organismal physiology. **b**, Schematic of engineered Yeast Artificial Chromosomes (YACs) with a *GAL1* promoter inserted to conditionally inactivate the centromere when cells are grown in galactose. **c-d**, Schematic for **c**, accumulation and **d**, curing of non-functional DNA in yeast cells. Centromeres were inactivated by growing cells on galactose for 5 – 7 cell divisions to accumulate multiple YACs in mother cells. Treated cells were plated on glucose-containing medium, where the centromere is re-activated, and YACs are stably propagated. Cells were plated at low density to isolate clonal cell lines carrying a defined number of YACs. Subsequent centromere inactivation on galactose and counter-selection allowed isolation of cell lines that lost all YACs (cured cells). **e**, DNA content analysis of 4 isolated clonal cell lines stably propagating different numbers of YAC-L (+nYAC-L). **f**, DNA content analysis of a cell line carrying multiple copies of YAC-L propagated in glucose-containing medium for the indicated time in non-selective conditions. **g**, DNA content analysis of 3 isolated colonies that were cured from YACs as described in **d**. **h**, Maximal growth rate of 10 strains carrying different numbers of accumulated YACs was measured on a plate reader, and doubling time is plotted against added DNA content measured by flow cytometry. **i**, Cell volume of the same strains as in **h** (showing 4 out of 10 strains) was measured on a Coulter counter (*left*), and mean cell volume is plotted against DNA content (*right)*. **j**, **k**, The YAC-bearing strains from **h** were cured from YACs as described in **d,** and growth rate **j** and mean cell volume **k** were measured and plotted against added DNA content.

We inactivated the YAC centromere for several consecutive generations, allowing the accumulation of multiple YAC copies (+nYACs) in mother cells. Subsequently, cells were transferred to glucose to re-activate centromeres and to allow stable YAC propagation. This procedure allowed us to isolate clones harboring up to one genome equivalent of YAC DNA that is stably maintained (**Fig. 1e-f, S1c**). These flow cytometry-based DNA content measurements were confirmed using DNA sequencing (Fig. **S1b**). Finally, YAC accumulation could be fully reversed by YAC centromere inactivation in cells containing multiple YACs and subsequent selection for cells that do not harbor any artificial chromosomes (**Fig. 1d, 1g**). We have therefore established a conditional and fully reversible genetic system to accumulate non-coding DNA in budding yeast cells.

### Accumulated non-coding DNA impairs cell fitness and leads to cell size increase

We next analyzed the phenotypic consequences of accumulating large amounts of non-coding DNA. Accumulation of human DNA led to a corresponding slowdown of cell growth (**Fig. 1h**), consistent with the idea that there is selective pressure against the expansion of non-coding DNA in the *S. cerevisiae* genome. Strikingly, we also observed that cell size increased proportionally to the amount of non-coding DNA (**Fig. 1i**). We found a similar coupling between cell size and DNA content when comparing the size of cells that varied by 20% in genome size because they contain different repeat copy numbers of the ribosomal DNA locus or in cells that accumulated high numbers of ectopic plasmids (**Fig. S1d-e**). These observations demonstrate that cell size is adjusted to total DNA content. Although the tight correlation between cell size and DNA content has been recognized for many decades ^13^, this is the first experimental demonstration that this coupling is direct and does not depend on the coding capacity of the DNA. Importantly, both the fitness penalty associated with DNA burden and cell size adjustment are fully reversible upon induced YAC loss (**Fig. 1j-k**), demonstrating that these phenotypes are direct consequences of harboring non-coding DNA.

### Non-coding DNA delays cell cycle progression

Because cell size and proliferation are tightly linked to cell cycle dynamics, we analyzed the cell cycle progression in cells burdened with non-coding DNA. Flow cytometry analysis revealed that the fraction of cells in G1-phase of the cell cycle increases with the amount of DNA burden (**Fig. 2a-b**). Consistent with excess DNA causing a delay in the duration of the G1-phase, cells harboring excess DNA release more slowly from a pheromone-induced cell cycle arrest in G1 (**Fig. 2c-d**). To gain further insight into how additional DNA affects cell cycle progression, we performed live-cell imaging of cells expressing Whi5-tdTomato, a cell cycle regulator that is exported from the nucleus at the G1 to S-phase transition ^22,23^ (**Fig. S2a**). Consistent with our observations from exponentially growing cultures, DNA burden led to a slight increase in G1-phase duration, although this difference was less pronounced in these growth conditions and did not reach statistical significance (**Fig. 2e, S2c**). In addition, excess DNA slows down progression through S/G2/M-phases (**Fig. 2d, 2f, S2c**). Thus, the presence of additional non-coding DNA slows down cell cycle progression at multiple stages.

**Figure 2.**
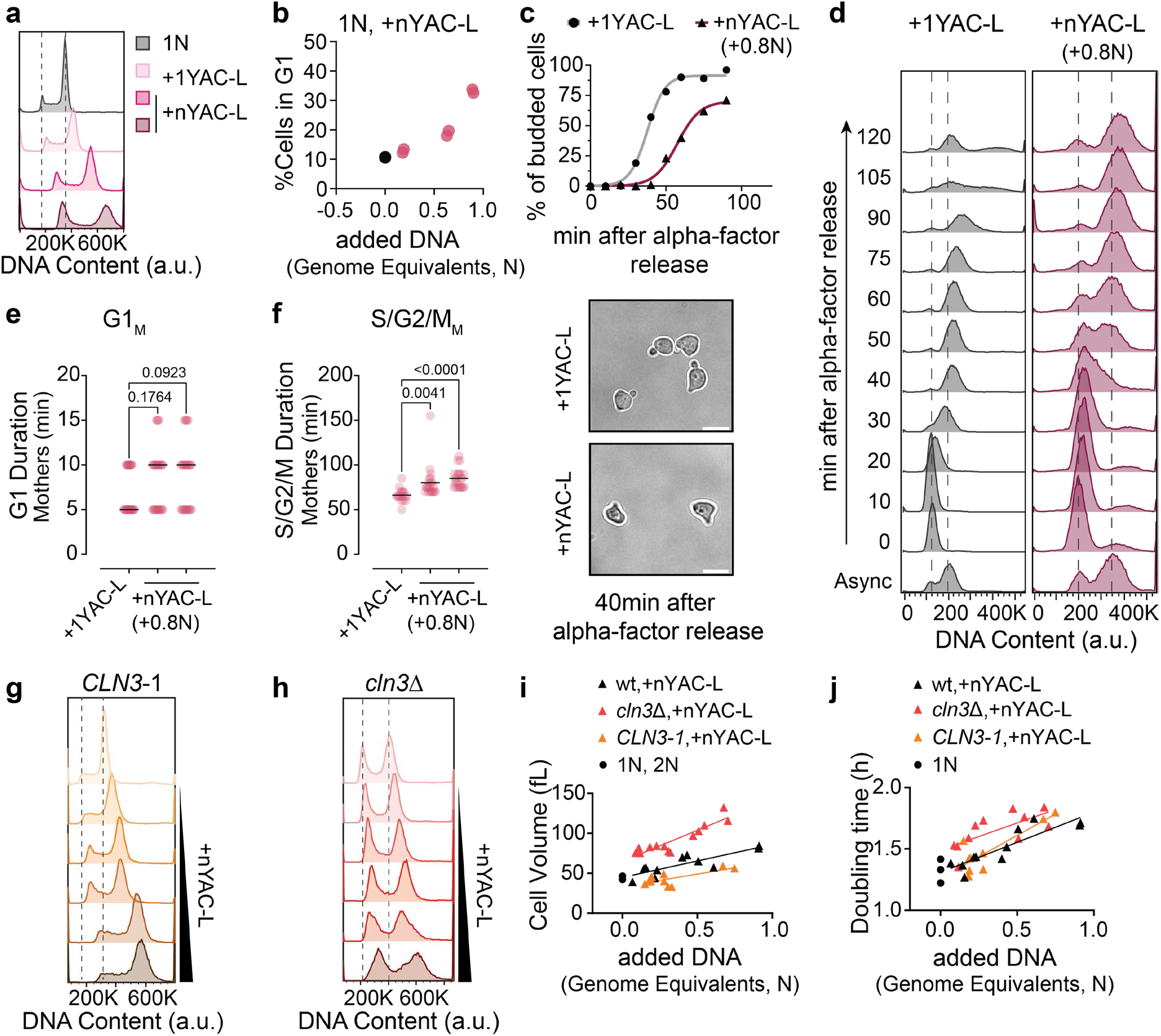
Non-coding DNA delays cell cycle progression. **a**, DNA content analysis of 3 isolated strains carrying different numbers of accumulated YACs. **b**, The fraction of cells in the G1-peak of the DNA content profile as shown in **a,** was measured and plotted against the added DNA content of the haploid control (1N, black), and isolated strains carrying different numbers of accumulated YACs (+nYAC-L, pink). 2 independent measurements are shown per cell line. **c**, **d**, Strains carrying one or multiple accumulated YACs were arrested in G1-phase using pheromone. The release from the cell cycle arrest upon pheromone washout was monitored by **c,** quantifying the fraction of budded cells and **d,** DNA content analysis. Representative images of cells after 40 min from G1 release. Scale bar, 5 µm. **e**, **f**, Live cell imaging of cells expressing Whi5-tdTomato carrying different numbers of YACs was performed, and the G1-phase duration (Whi5 nuclear) and S/G2/M duration (Whi5 cytoplasmic) were measured in mother cells (median, black line). >30 cells were analyzed per condition. Adjusted p values were calculated by Kruskal-Wallis test followed by Dunn’s multiple comparisons correction. The data from **e**, **f** is generated from two independently isolated YAC-bearing strains. One representative dataset from 2 independent measurements is shown. **g**, **h**, DNA content analysis of **g**, *CLN3-1* and **h**, *cln3*Δ strains carrying different numbers of YACs. **i**, Mean cell volume of wild-type, *CLN3-1*, and *cln3*Δ strains carrying different numbers of YACs plotted against added DNA content. **j**, Growth rate of wild-type, *CLN3-1*, and *cln3*Δ strains carrying different numbers of YACs was measured, and doubling time is plotted against added DNA content. The data from **g**-**j** is generated from at least two independently isolated YAC-bearing strains per genotype and from ≥ 3 independent measurements.

### G1 delay is not required to adjust cell size to DNA content

In budding yeast, cell size is primarily adjusted during the G1 phase ^24^. Consistent with this model, cells harboring large amounts of non-coding DNA delay cell cycle entry and reach a larger threshold cell size before entering cell division (**Fig. S2d**), showing characteristics of both a “sizer” and an “adder” size control mechanism ^25^ (**Fig. S2e-f**). These observations suggest that DNA content may be measured during G1 phase to adjust cell size accordingly. We therefore determined whether the observed G1 delay is required for the scaling of cell size to DNA content. For this purpose, we analyzed mutant strains that either accelerate or delay progression through G1 because of mutations in the gene encoding the G1 cyclin Cln3 ^26^. Cells expressing the stabilized *CLN3-1* allele have extremely short G1-phases that are often not detectable by flow cytometry, while deleting *CLN3* increases the fraction of cells in G1 (**Fig. 2g-h**). Deleting *CLN3* increased cell size, as previously reported, and it also slightly increased the DNA-dependent cell size adjustment (**Fig. 2i**), indicating that Cln3 contributes to overcoming the DNA-dependent G1-delay. However, stabilizing Cln3 did not abolish the correlation between cell size and DNA content, demonstrating that the increase in G1 duration caused by excess DNA burden is not necessary to couple cell size to DNA content. While deleting *CLN3* mildly affected the correlation between DNA content and cell size, the fitness penalty associated with the accumulation of non-coding DNA was not affected by either mutation in *CLN3* (**Fig. 2j**). This observation suggests that the fitness defect caused by non-coding DNA is independent of the DNA induced cell size increase.

### Human DNA is inefficiently replicated in yeast

Apart from delaying the duration of G1-phase, excess DNA burden also slows down cell cycle progression through S and G2/M phases (**Fig. 2f, S2c**). A possible explanation for this observation is that increasing DNA content might prolong the time it takes to replicate the genome, as excess DNA might titrate limiting replication factors and prolong the S-phase accordingly. To test this hypothesis, we determined how the presence of additional DNA affects genome replication using Sort-seq ^27^. For this purpose, we isolated S-phase and G2-cells using fluorescence-activated cell sorting (FACS) and determined the relative DNA copy number of these sub-populations using DNA sequencing. This analysis allows accurate and robust identification of early- and late-replicating genome regions, which can be used to approximate replication timing dynamics ^28,29^.

This DNA replication analysis revealed that the replication timing of endogenous yeast chromosomes is mildly affected by the accumulation of large amounts of exogenous DNA (**Fig. 3a, S3a-c**). In select genomic regions, replication peaks are lower and broader, indicating local differences in origin firing and fork progression speed. However, overall, the effects of excess DNA burden on the replication program of yeast chromosomes are modest. In contrast, human DNA sequences are replicated less efficiently in *S. cerevisiae* than endogenous chromosomes (**Fig. 3b**). Importantly, the relative replication timing of the exogenous DNA does not progressively worsen with increasing DNA burden (**Fig. 3b**), suggesting that the late replication of human DNA in yeast is not a consequence of replication factors becoming limiting.

**Figure 3.**
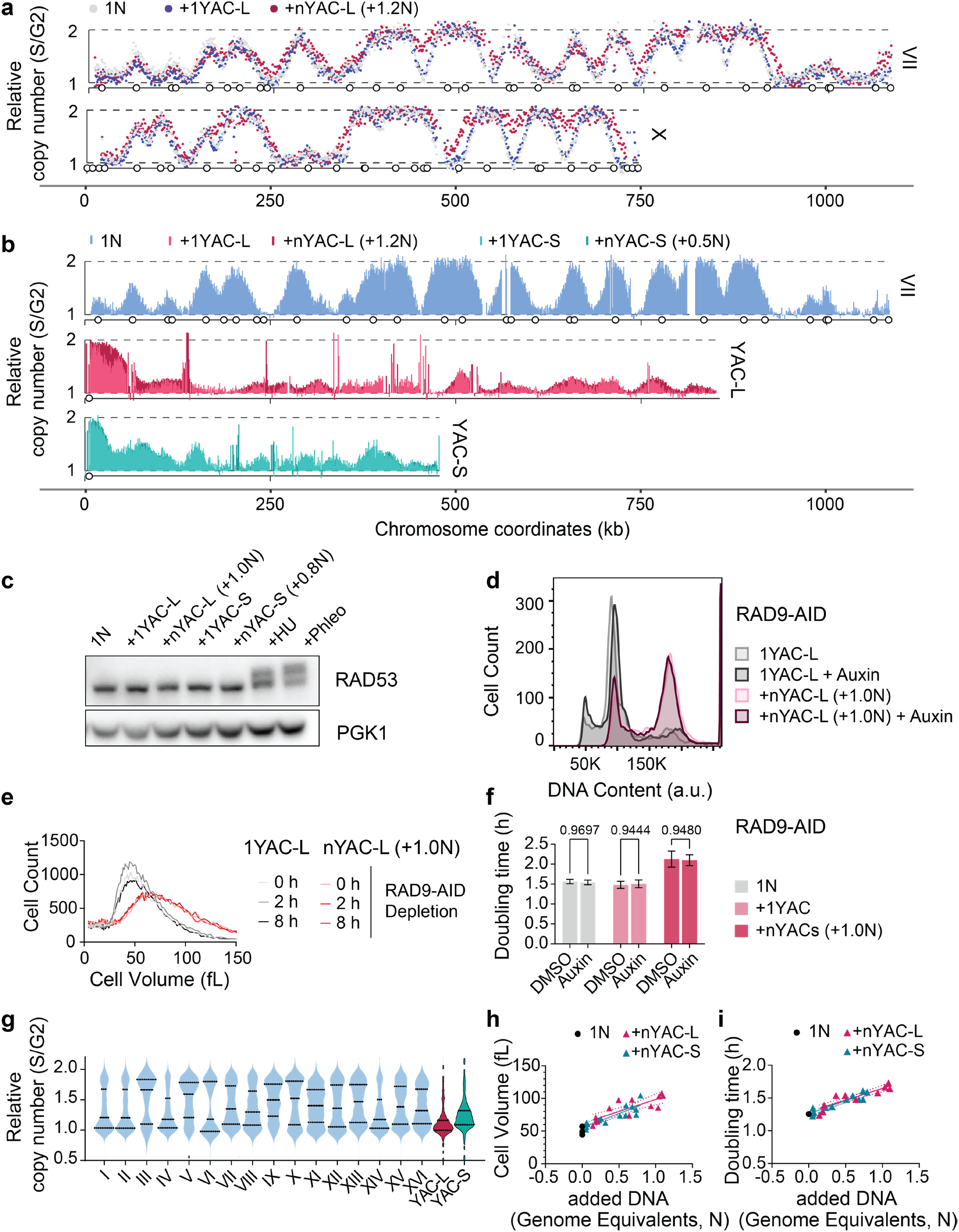
Increasing genome size has little effect on the yeast genome replication program. **a**, S-phase and G2-phase cells were sorted by flow cytometry and sequenced to determine the relative copy number ratios (S/G2) per genomic coordinate during S-phase (Sort-seq). The replication profiles of *Saccharomyces cerevisiae* chrVII and chrX are shown for a haploid control (1N: NF=1.425), a strain with one YAC-L (1YAC-L: NF=1.403), or multiple YAC-L copies (nYAC-L: NF=1.388). Circles indicate annotated yeast origins of replication (ARS). Raw data were normalized to scale values between 1 and 2. The normalization factors (NF) are indicated in brackets. **b**, Sort-seq replication profiles of *S. cerevisiae* chrVII from haploid control cells (1N: 1.456) and of the artificial chromosomes YAC-L (1YAC-L: NF=1.403, nYAC-L: NF=1.338) and YAC-S (1YAC-S: NF=1.451, nYAC-S: NF=1.366) in strains carrying either one or multiple YACs. Circles indicate annotated yeast origins of replication (ARS). **c**, Western blot showing total RAD53 in lysates of exponentially growing cultures of strains carrying different numbers of the indicated accumulated YACs. Wildtype cells treated with hydroxyurea (50 mM) and phleomycin (50 µg/ml) for 30 minutes are shown as positive controls. Representative of 2 biological replicates. **d**, DNA content analysis in cells carrying one or multiple YACs before and after depletion of Rad9 using the auxin-inducible degron (AID) system for 8 hours (100 µM IAA). Analysis of 2 biological replicates. **e**, Coulter counter cell volume measurements in cells carrying one or multiple YACs before and after depletion of Rad9-AID. Analysis of 2 biological replicates. **f**, Growth rates of cells carrying one or multiple YACs and expressing the Rad9-AID allele with and without depletion of Rad9-AID by addition of Auxin. Analysis of 3 biological replicates. Adjusted p values were calculated by two-way ANOVA followed by Šidák’s multiple comparisons test. **g**, Distribution of relative copy number ratio in sorted S versus G2 cells values per genomic coordinate of each *S. cerevisiae* chromosome and YAC-L and YAC-S. **h**, Mean cell volume of strains carrying different numbers of either YAC-L or YAC-S plotted against added DNA content. Analysis of 2 isolated colonies with different amounts of accumulated non-functional DNA from 3 independent experiments. **i**, Growth rate of strains carrying different numbers of either YAC-L or YAC-S was measured, and the doubling time is plotted against added DNA content. Analysis of 2 isolated colonies with different amounts of accumulated non-functional DNA from 3 independent experiments.

Of note, large yeast chromosomes also harbor late-replicating regions that are comparable in size to the 850-kb YAC-L (**Fig. S3c**). In particular, the megabase-sized rDNA array and sub-telomeric regions replicate late and only finish replication after cells have entered mitosis ^30,31^. Together, these observations raise the interesting possibility that expansion of non-functional DNA is prevented because it interferes with effective genome replication and slows down cell proliferation.

### Replication stress is not responsible for loss of cellular fitness upon non-coding DNA accumulation

Next, we set out to determine whether the accumulation of non-coding DNA causes replication stress. For this purpose, we measured cell growth in the presence of a sublethal dose of hydroxyurea (HU), a drug that blocks deoxynucleotide synthesis and that is particularly toxic for cells experiencing DNA replication stress ^32^ (**Fig. S3d**). Indeed, cells bearing excess DNA burden grow slightly slower in the presence of HU, suggesting that accumulation of late-replicating DNA might lead to a basal level of replication stress. To directly address this possibility, we analyzed the phosphorylation status of Rad53, the effector kinase that is activated by DNA damage and replication stress ^33^. Rad53 phosphorylation was found indistinguishable between cells carrying no and multiple YACs (**Fig. 3c**), suggesting that YAC-bearing cells do not have elevated basal levels of DNA replication stress. To further probe whether DNA damage signaling delays cell cycle progression in cells carrying excess DNA burden, we depleted the checkpoint adaptor protein Rad9 using the auxin-induced degron system ^34^ (**Fig. S3e**). Rad9 depletion did not alter the DNA content or cell cycle profile of cells bearing an excess DNA burden (**Fig. 3d**), demonstrating that successful YAC replication does not rely on checkpoint activation. Similarly, Rad9 depletion did not alter cell size within 8 hours of checkpoint inactivation, and it did not affect the fitness penalty incurred by DNA burden (**Fig. 3e-f**). These results show that the phenotypes associated with excess non-coding DNA are independent of DNA damage signaling.

### Cell size adjustment and fitness defects are not caused by the slow replication of YACs

To directly test whether late replication of exogenous DNA is, at least partially, responsible for the phenotypes caused by the presence of non-coding DNA, we sought to improve the replication efficiency of the exogenous DNA. Insertion of the well-described early firing replication origin *ARS1* ^35^ only slightly improved YAC replication, indicating that the DNA context, rather than the origin sequence, determines the timing of origin firing (**Fig. S3f-g**).

The only early firing replication origin on the YAC is located close to the centromere at the very beginning of the long chromosome arm (**Fig. 3b**). We therefore reasoned that we could improve the fraction of early versus late replicating DNA by using the shorter 450 kb YAC-S, which contains different human DNA regions but has an overall similar sequence composition to the larger 850kb YAC-L (**Fig. S1a**). Indeed, the fraction of early replicating regions is significantly increased on YAC-S compared to YAC-L (**Fig. 3g**). Notably, improving YAC replication did not alter the correlations between DNA content, cell size, and cell fitness (**Fig. 3h-i**), demonstrating that late replication of exogenous DNA is not the primary cause of impaired cell fitness or of the coupling of cell size to DNA content. In addition, the comparison between these two different YACs shows that the cell size to DNA coupling is not mediated by the number of centromeres or by other sequence-specific elements. In agreement with this conclusion, expansion of non-coding DNA without changing the number of centromeres leads to the same cell size and fitness phenotypes ^36^. Together, our results show that the accumulation of non-coding DNA impairs cell fitness and leads to an increase in cell size that is independent of the cell size and DNA damage checkpoints.

### Non-coding DNA titrates RNA polymerase away from endogenous chromosomes

Previous work showed that the transcription machinery in budding yeast engages with random DNA sequences ^37,38^. We therefore wondered whether accumulation of non-coding DNA might affect the production of endogenous transcripts. To determine how DNA burden affects transcription, we performed spike-in normalized RNA sequencing of cells bearing excess human DNA. As previously reported ^37,38^, we observed pervasive transcription of human DNA in budding yeast. In cells carrying one genome equivalent of non-coding DNA, reads from human DNA made up 6% of total RNA reads, excluding ribosomal RNA (**Fig. 4a**). Importantly, work by Lu et al. shows that transcripts from YAC DNA are rarely translated into proteins ^36^, arguing against the possibility that the fitness burden associated with exogenous DNA is mediated by YAC-derived peptides.

**Figure 4.**
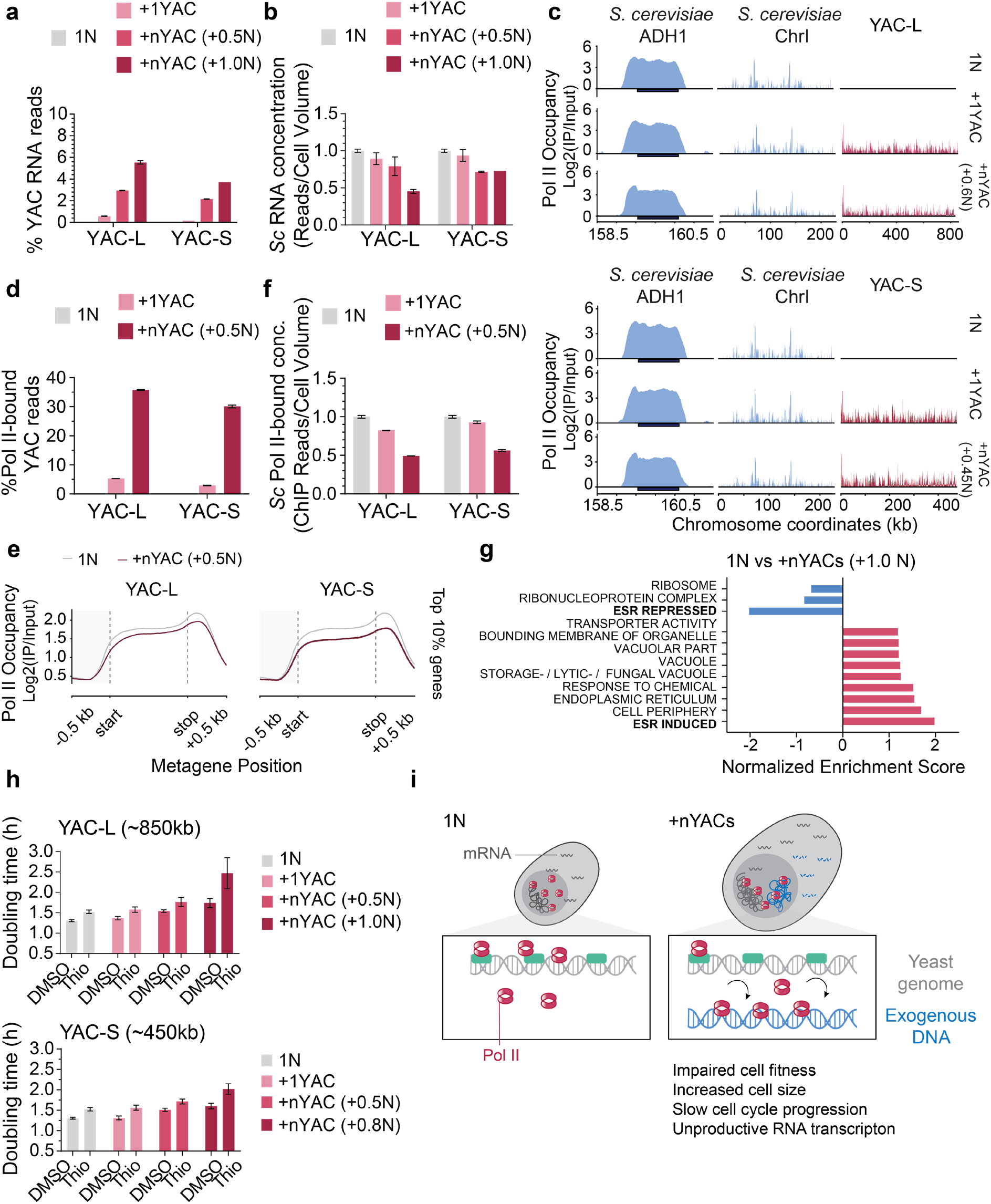
Non-functional DNA titrates transcription machinery components from essential genes. **a**, Spike-in total RNA sequencing was performed in cells carrying none or different numbers of the indicated YAC. Percentage of RNA reads, excluding rRNA reads, from exogenous DNA is shown. The mean and range (error bars) of 2 biological replicates are shown. **b**, Concentration of *S. cerevisiae* reads was determined by normalizing the *S. cerevisiae* reads to the spike-in *C. albicans* reads and dividing this number by the mean cell volume of the respective strain. The values are normalized to the haploid control (1N) for comparison. The mean and range (error bars) of 2 biological replicates are shown. **c**, Genome-wide Pol II occupancy profiles. Genome browser tracks showing the log2 enrichment ratio of Pol II ChIP over Input (log2 [ChIP/Input]) across genomic coordinates. Signals from 2 pooled biological replicates were independently normalized to exogenous *C. glabrata* spike-in reads before ratio calculation. See *Materials and Methods* for detailed normalization procedures. In **c**-**e** and **g** the amount of added YAC DNA was estimated based on the ratio of total YAC reads to Sc reads in the ChIP Input sample. **d**, Bar chart representing the percentage of total Pol II ChIP-seq reads mapping to the YAC sequence of the indicated strains. See *Materials and Methods* for detailed normalization procedures. The mean and range (error bars) of 2 biological replicates are shown. **e**, Metagene profiles showing Pol II occupancy across gene bodies (scaled to 1 kb with 500 bp flanking regions) for the top 10% most heavily occupied genes in the baseline 1N control strain. See *Materials and Methods* for profiling details. Signal from 2 pooled biological replicates. **f**, The cellular concentration of Pol II bound to the *S. cerevisiae* genome was calculated by dividing the spike-in normalized sum of Sc ChIP reads by the respective cellular volume of each strain. The values are normalized to the haploid control (1N) for comparison. The mean and range (error bars) of 2 biological replicates are shown. **g**, Gene set enrichment analysis (GSEA) of the RNA-seq experiment shown in **a**-**b**, comparing gene expression in haploid control strains with strains containing 1-genome-equivalent exogenous DNA. Data for YAC-L and YAC-S bearing strains were combined from 2 biological replicates for this analysis. Displaying the top-ranked gene sets by normalized enrichment scores (NES). Gene sets displaying increased expression in DNA-burdened cells are shown in red, downregulated gene sets in blue. **h**, Growth of strains carrying different YAC numbers was measured in the presence or absence of the RNA Polymerase II inhibitor Thiolutin (1.6 mg/ml). Each bar represents the mean and standard deviation of 2 independent clones measured in 3 independent experiments. **i**, Schematic of the consequences of excess DNA in budding yeast cells. Accumulation of non-coding DNA leads to impaired proliferation, cell enlargement, and cell cycle slowdown. Specifically, sequestration of transcription machinery away from yeast genes can explain the loss of competitive fitness in the presence of an excess of non-coding DNA.

Instead, in cells with a high non-coding DNA burden, the increased number of exogenous transcripts coincided with decreasing yeast transcript concentrations (**Fig. 4b**), suggesting that non-coding DNA interferes with transcription of endogenous genes. To directly determine binding of RNA Polymerase II (Pol II) to endogenous and exogenous DNA, we performed a calibrated spike-in ChIP-sequencing experiment against the Pol II subunit Rpb1 ^39,40^. This analysis showed that RNA polymerase binds to exogenous DNA nearly as densely as to yeast chromosomes containing functional elements (**Fig. 4c, S4a**). Consequently, in a strain burdened with 0.5 genome equivalents of exogenous DNA, ∼35% of Pol II was bound to exogenous DNA (**Fig. 4d, S4b**). As YAC transcripts made up only 6% of total transcripts in the same strains, these data indicate that either the transcription of human DNA is less efficient or that the transcripts originating from YAC DNA are less stable than endogenous mRNAs, or both.

Previous work showed that the amount of RNA polymerase bound to chromosomes increases as cells grow larger, reflecting the increasing mRNA synthesis rate with cell size ^40^. In contrast, Pol II occupancy on gene bodies decreases in DNA-burdened cells (**Fig. 4e**), even though these cells are larger than cells without DNA burden. To take cell size into account, we divided the number of Pol II-bound reads mapping to yeast DNA (**Fig. S4b**) by the mean cell volume of the respective strain to calculate the cellular concentration of Pol II engaged in productive transcription. When compared to haploid control cells, this number is reduced by 44-50% in cells burdened with 0.5 genome equivalents of exogenous DNA (**Fig. 4f**). Titration of RNA polymerase to non-coding DNA therefore prevents cells from producing sufficient endogenous mRNAs to support their size.

Interestingly, sequestration of the transcription machinery from endogenous chromosomes most severely affected highly expressed genes, which are enriched for essential housekeeping genes, such as ribosomal protein genes (**Fig**. **4g**). This reduced expression of ribosomal genes is part of a stereotypical transcriptional signature that is associated with slow growth ^41^ (**Fig. S4c)**. We reasoned that failure to transcribe ribosomal genes at sufficiently high rates could explain the fitness penalty associated with excess DNA burden. In agreement with this, previous work showed that transcription can become limiting for cell growth under certain conditions ^42–44^. To test whether transcription becomes limiting for growth in DNA-burdened cells, we treated cells with sub-lethal doses of the Pol II inhibitor Thiolutin ^45^. Cells carrying excess non-coding DNA became hypersensitive to transcription inhibition (**Fig. 4h**), demonstrating that the transcription machinery becomes limiting for cell growth in DNA-burdened cells. Together, our results show that non-coding DNA impairs cell fitness in budding yeast by sequestering the limiting transcription machinery.

## Discussion

How the accumulation of non-coding DNA impacts cell physiology is a key question in understanding the evolution of genomes. Here, we developed a genetic tool that allows characterization of the consequences of increasing DNA content without altering gene dosage. We find that cell size is coupled to DNA content and that accumulation of non-coding DNA impairs cell proliferation by titrating the limiting transcriptional machinery away from endogenous chromosomes.

Maintaining a constant DNA:cytoplasm ratio is critical to prevent cellular dysfunction. At critically low ratios, DNA limits transcription, causing cytoplasmic dilution and severe genome homeostasis defects ^43,46,47^. Conversely, an acute increase in the DNA:cytoplasm ratio following polyploidization triggers severe DNA replication stress ^48^. How cells couple cell size to DNA content has, however, remained unclear. Because the expression level of important cell cycle regulators is a key determinant of cell size ^49–51^, gene copy number variation has been hypothesized to link cell size to ploidy.

Here, we provide the first experimental evidence that the coupling of cell size to DNA content occurs even if the additional DNA does not encode functional proteins. Our work demonstrates that this coupling is direct, reversible, and occurs within one or a few generations. The correlation of cell size and genome content across species therefore does not rely on co-evolution but rather on direct coupling. Mechanistically, this coupling does not require the canonical G1/S checkpoints and DNA damage signaling networks, suggesting that it relies on fundamental metabolic or biophysical processes. While it remains unclear which cellular processes these are, it is now possible to address this long-standing question experimentally using our developed strategy to modulate DNA content.

Even though whole-genome duplications (polyploidy) and chromosomal imbalances (aneuploidy) are hallmarks of tumor genomes and were the focus of intense studies ^10–12^, the physical burden of excess DNA has been widely overlooked. Introduction of exogenous DNA represents a relatively small increase in absolute biomass, with DNA content only corresponding to 1-2% of cell biomass ^52^ and histones representing less than 1% of all proteins ^53^. The energetic cost of producing twice the amount of chromatin is therefore relatively small and unlikely to explain the 30% increase in doubling time upon addition of one genome equivalent of non-coding DNA. Instead, our analysis revealed that RNA polymerase is recruited to exogenous DNA as efficiently as to yeast DNA, consistent with previous observations ^37,38^. We show that this pervasive transcription of non-coding DNA occurs at the expense of the production of endogenous transcripts, which are less concentrated in DNA-burdened cells. Importantly, we find that these cells become hypersensitive to transcription inhibitors, indicating that transcription is limiting cell growth in DNA-burdened cells.

*S. cerevisiae* has lost key components of the constitutive heterochromatin and RNA interference pathways ^54^, and we show here that exogenous DNA sequences are bound as efficiently by RNA polymerase as endogenous DNA. In contrast, random DNA sequences are repressed efficiently in vertebrate cells and likely in other species with intact heterochromatin pathways ^37,38^. We therefore propose that the inability to repress non-coding DNA increases the fitness cost of non-functional DNA in yeast and prevents the excessive expansion of non-coding DNA elements that occurred in complex eukaryotes.

## Methods

### Yeast strains and growth conditions

All yeast strains used in this study are listed in Supplementary Table 1. All strains are derived from W303 background. Yeast strains containing one YAC copy of different sequence identity and length were previously described in detail ^55^, and are based on the vector pYAC4 ^56^. All original yeast strains containing YAC sequences are listed in Supplementary Table 2. Yeast artificial chromosomes are linear DNA fragments containing large human DNA fragments, an origin of replication (*ARS1*), a centromere (*CEN4*), and telomeres. The YACs used in this study contain two auxotroph selection marker genes (*URA3, TRP1)*, allowing us to select for cells with at least one artificial chromosome copy and to select for cells that have lost all YACs using 5-Fluoroorotic Acid (5-FOA, 1 mg/mL). Protein fusions and gene deletions were generated using PCR-based integration methods ^57,58^.

Yeast media were prepared according to standard laboratory practices ^59^. Liquid cultures were grown in shaking flasks at 220 r.p.m. in YPD (1% yeast extract, 2% peptone, and 2% dextrose) medium. For imaging experiments, cells were grown in synthetic dextrose (SD) medium (0.17% Yeast nitrogen base without amino acids, 2% carbon source, 0.5% NH_4_ sulfate, and 1x amino acids mix). Cells were grown at 30 °C except for **Fig. 3d-f** where cells were grown at 25 °C, as the *RAD9-AID* allele is hypomorphic at higher temperatures.

### YAC accumulation

To accumulate non-coding DNA in haploid yeast cells, we triggered the missegregation of YACs by inactivating their conditional centromere. Yeast cultures harboring a YAC containing a *GAL1* promoter adjacent to its centromere were grown in Yeast extract-peptone (YP) 2% raffinose for 24 hours and subsequently grown in YP 2% raffinose 1% galactose for 5 to 7 cell generations. 200 – 2000 yeast cells were plated on SD-URA plates to isolate single clonal populations, each carrying a defined YAC copy number. YAC accumulation was quantified for each experiment by nucleic acid staining and flow cytometry (see below).

### YAC curing

To exclude that the observed phenotypes were caused by other factors than the acquired non-coding DNA, YAC loss was induced by missegregation and subsequent counter-selection. For this process, strains bearing one or multiple YACs were streaked on YP plates with 2% raffinose and 1% galactose to trigger YAC missegregation. Cells were subsequently streaked on SD plates containing 5-Fluoroorotic Acid (5-FOA, 1 mg/mL) to select for cells that have lost all YAC copies.

### G1-phase cell cycle arrest and release

To block yeast cells in G1, yeast strains were grown to an O.D._600_ of 0.25 in YPD and treated with 5 µg/mL α-factor (GeneScript) for a total of 2.5 hours at 30 °C. An additional 2.5 µg/mL α-factor was added after 1 hour of treatment. For the G1-release, yeast cultures were collected on a 0.45 µm pore size filter membrane (Millipore) by vacuum-filtration and washed twice with media. Collected cells were resuspended in YPD and grown at 30°C. Samples were taken at regular intervals up to 120 minutes after release to determine budding index and for DNA content analysis. To determine budding index, cells were fixed in 4% formaldehyde for 5 minutes, washed twice in Phosphate-buffered saline (PBS), and briefly sonicated to break up cell clumps (2 cycles, 20%, 2 seconds on a BANDELIN Ultrasonic Homogeniser HD 4100). The fraction of budded cells was determined from > 30 cells per condition using brightfield illumination on a Nikon Eclipse Ti-E inverted microscope using a 60x/NA 1.40 oil objective.

### Cell growth and size measurement

For growth curve analysis, overnight yeast cultures were diluted to an OD_600_ of 0.02 in 48-well plates with the indicated media. Cell growth at 30 °C was monitored by absorbance at 600 nm wavelength every 5 minutes for 48 hours on the plate reader SPECTROstar Nano (BMG Labtech).

To measure cell volume, we briefly sonicated logarithmically growing yeast cells, diluted in 10 mL Isotone II (Beckman), and measured on a Coulter Counter (Multisizer 4e, Beckman).

### DNA content analysis

To quantify the amount of accumulated non-coding DNA, we measured the DNA content by flow cytometry ^60^. In short, we fixed one volume of yeast cultures with two volumes of 100% ethanol and incubated at -20 °C overnight. Fixed cells were subsequently pelleted at 4 °C and washed twice with 50 mM Sodium Citrate (pH 7.2) with a 10-minute incubation at room temperature between washes. Cell pellets were then resuspended in 50 mM Sodium Citrate (pH 7.2) containing 2.5 µM SYTOX Green Nucleic Acid stain (Invitrogen) and 100 µg/ml RNase A (Sigma-Aldrich), and incubated for one hour at 37 °C, shaking at 440 r.p.m, in the dark. Samples were later digested with 0.4 mg/mL Proteinase K (Panreac AppliChem) for one hour at 55°C, shaking at 440 r.p.m, in the dark. To break up cell clumps, stained yeast cells were sonicated (2 cycles, 20%, 2 seconds on a BANDELIN Ultrasonic Homogeniser HD 4100). DNA content was measured using a 488 nm laser, 530/30 optical setup on the cell analyzer LSRFortessa (BD), or 488 nm laser, 533/30 optical setup on the cell analyzer Accuri C6 Plus (BD), collecting at least 10,000 events per sample.

To estimate the amount of accumulated YAC DNA, DNA content analysis results were analyzed using FlowJo Software (v10.8, 2023). The univariate model Watson Pragmatic algorithm ^62^ was used to determine the G1 peak signal for each strain. The signal was then normalized to the G1 peak of the haploid control *wild type* or the respective *mutant* strain to determine the amount of added non-coding DNA. For experiments in **Fig. 2g-j**, where the G1 peak was not clearly defined, the G2 peak was used for genome content normalization. YAC DNA content for the ChIP-seq experiment (**Fig. 4c-f**, **4g**, **S4a-b**) was estimated using the total read counts of the Input samples and calculated by dividing the total YAC reads by the sum of *S. cerevisiae* reads. The DNA content determined by this method is roughly 30% lower than DNA content determined by flow cytometry, likely because of the relatively poor sequencing coverage of the YAC DNA that might be caused by the degree of repetitive sequences (**Fig. S3a**).

### Pulsed-field gel electrophoresis

To independently verify the presence of the YACs in yeast strains, we resolved and visualized chromosomes using PFGE as described in the manual for the Bio-Rad CHEF-DR-III. In short, yeast cultures were grown to exponential growth phase in YPD, and 1.25 x 10^8^ cells were collected by centrifugation and washed in ice-cold 50 mM Ethylenediaminetetraacetic acid (EDTA). Cells were resuspended and washed once in cell suspension buffer (10 mM Tris, 50 mM EDTA, and 2 mM NaCl). Cell pellets were resuspended in buffer containing 52 U/mL Zymolyase (Amsbio) and 2% molten CleanCut agarose (Bio-Rad). The cell mixture was then pipetted into a plug mold (Bio-Rad). Plugs were cooled down at 4 °C for 10 minutes and pushed out into microcentrifuge tubes with 0.5 mL of buffer containing 30 U/mL Zymolyase. The plugs were incubated at 37 °C for 2 hours and transferred to new microcentrifuge tubes containing 0.75 mL proteinase K buffer (10 mM Tris, 100 mM EDTA, and 0.5% SDS) with 0.8 mg/mL proteinase K (PanReac AppliChem). The plugs were then incubated at 50 °C overnight. Subsequently, they were transferred into tubes containing 0.5× Tris base-Borate-EDTA buffer (TBE) (45 mM Tris base, 45 mM boric acid, and 1 mM EDTA, pH 8.3) and equilibrated for 45 minutes. Plugs were then placed into the wells of the running gel made up of 1% Agarose (Pulsed-Field-Certified Agarose, Bio-Rad) in 0.5 × TBE. Gels were run on a CHEF-DR-III Pulsed-Field Gel Electrophoresis System (Bio-Rad), at 6 V cm−1, 120°, switch time 60–120 s, for 24 hours at 14 °C. The gel was stained with GelRed (Biotium) for visualization and imaged on GelDoc Molecular Imager XR+.

### Live-cell microscopy

To monitor the cell cycle progression of YAC-containing strains, all cells were grown in SD 2% glucose overnight at 30°C. Cultures were diluted to an OD_600_ of 0.05 and grown for 3 to 4 hours before imaging. Cells were plated in 8-well µ-slide chambers (Ibidi), previously incubated with 2 mg/mL Concanavalin A (ConA) (Sigma) for 15 minutes and washed three times with SD 2% Glucose. Live cell imaging was performed on a Nikon Eclipse Ti-E inverted microscope using a 60x/NA 1.40 oil objective. mCherry fluorescence was imaged using a 575/25 nm excitation filter (Lumencor) and a 641/75 nm emission filter (AHF Analysentechnik). GFP fluorescence was imaged using a 470/40 nm excitation filter (F49-470) and a 520/35 nm emission filter (AHF Analysentechnik). Images were acquired with a Hamamatsu ORCA Flash 4.0 camera controlled by the NIS software (Nikon). Three to four fields of view were collected per condition.

#### Image analysis

To assess cell cycle phase progression, cells were followed in the bright light channel, and fluorescent images were acquired every 5 minutes. Image processing (cropping, Z-projection, tracing, etc.) and manual analysis were performed using Image J (version 2.16.0) ^63^. Only cells in focus were considered for the analysis, and at least 30 mother and 30 daughter cells per condition were assessed for the duration of one cell cycle. G1 cell cycle phase duration was determined from the time of nuclear Whi5 appearance to Whi5 export outside the nucleus, and S/G2/M combined cell cycle phase duration was determined from Whi5 nuclear export to the next Whi5 nuclear import. In mother cells where Whi5 nuclear import was not clearly detectable due to a very short G1-phase, Whi5 nuclear re-import in the corresponding daughter cell was used to approximate mitotic exit. Cell volume was determined by measuring the major and minor axes of an ellipse traced over cells (assuming yeast cell volume is an oblate spheroid shape). Area measurements were taken at the time of Whi5 nuclear export for critical cell volume estimation and at the time of Whi5 nuclear import in daughter cells for birth cell volume estimation.

### DNA replication timing using S-phase sequencing

To determine DNA replication timing in YAC-containing strains, we used a sequencing-based method, Sort-Seq ^27^. This method measures the relative copy number per genome coordinate of sorted S-phase and G2-phase cells. Yeast cultures were grown overnight and diluted to 0.05 O.D._600_ into 25 ml YPD medium and grown at 30 °C to a density of 0.5 – 0.9 O.D._600_. Cells were collected by centrifugation at 2,000 g for 5 minutes at 20 °C and washed twice with distilled water. Yeast cells were then fixed in 70% ethanol and incubated at 4 °C overnight. Fixed cells were washed twice with 50 mM sodium citrate (pH 7.2) and resuspended in 1 mL 50 mM sodium citrate and treated with 2 mg/mL RNase A for 1 hour at 37 °C, followed by a 1.48 mg/mL Proteinase K digestion for 1 hour at 55 °C. Yeast cells were stained with 10 µM SYTOX Green Nucleic Acid stain (Invitrogen) in 50 mM sodium citrate containing sodium azide (0.1%) and incubated at 4 °C overnight. Stained cells were sonicated (2 cycles, 20%, 3 seconds on a BANDELIN Ultrasonic Homogeniser HD 4100) and filtered through a 50 µm filter (CellTricks) to get rid of cell clumps. Stained cells were then sorted on a FACSAriaIII (BD) by DNA content using a 488 nm laser with a 530/30 filter and gating for S-phase and late G2-phase cells from singletgated cells, excluding cell debris.

We collected 7.5 million cells from S-phase and G2 sub-populations. Additionally, we collected 300,000 single cells per strain to assess sorting purity. The purity of the sorted cell fractions was confirmed later by nucleic acid re-staining, measured on the cell analyzer LSRFortessa (BD). Sorted cells were flocculated by adding 1/3 volume of 100% ethanol and pelleted by centrifugation at 20,000 g for 10 minutes. Cell pellets were resuspended in 100% ethanol and kept at 4 °C to preserve the sample until DNA extraction.

For DNA extraction, cells were pelleted and resuspended in 1.2 M Sorbitol, 0.2 M Tris-HCl (pH 8.0), 0.02 M EDTA (pH 8.0), and incubated with 1 mg/mL Zymolyase and 0.001% β-mercaptoethanol at 37 °C for 30 minutes. After digestion, samples were supplemented with 0.42% SDS, 85 mM NaCl, 83.3 µg/mL RNase A, and 0.33 mg/mL Proteinase K and incubated for 2 hours at 55 °C. Samples were then mixed with an equal volume of phenol:chloroform:isoamyl alcohol (25:24:1, pH 7.5-8.0, Carl Roth), and the phases were separated by centrifugation (16 000g for 5 min). The aqueous phase was transferred to a new tube and mixed first with an equal volume of phenol: chloroform: isoamyl alcohol, then with chloroform. To precipitate the DNA, the aqueous phase was mixed with a 2:1 volume of 100% ethanol and incubated at -20 °C overnight. The resulting DNA pellet was washed once with 70% ethanol, air-dried, and dissolved in 100 µl TE buffer (10 mM Tris pH 8.0, 1 mM EDTA). DNA samples were submitted for NEB Next Ultra II (New England Biolabs) library preparation and sequenced by 150 bp paired-end Illumina sequencing.

Reads were pre-processed to remove low-quality sequences and mapped using bowtie2 ^64^ (version 2.5.2) and subsequently mapped with a modified localMapper pipeline (https://github.com/DNAReplicationLab/localMapper/). Read counts were combined into 1-kb genomic bins, and replication profiles were generated by dividing S-phase reads by G2-phase reads using the Repliscope R package (https://cran.r-project.org/web/packages/Repliscope/). The S/G2 ratios were normalized to fit all ratio values to a biologically relevant scale ranging from 1 to 2. The normalization factor used is indicated for each sample in the figure legends.

### Spike-in normalized RNA-seq analysis

To perform transcriptome analysis on total cellular RNA from *Saccharomyces cerevisiae* YAC-containing strains, cells were grown in YPD to exponential phase (OD_600_ = 0.3). As cell size varied for different strains, the exact cell number was determined on a Multisizer 4e Coulter counter. An equal number (6.0 x 10^7^) of cells per strain was collected by centrifugation. Cells were washed once with DEPC-treated water, and pellets were snap-frozen in liquid nitrogen. Cell pellets were resuspended in 400 µL TES buffer (10 mM Tris pH 7.5, 10 mM EDTA, 0.5% SDS) combined with previously collected *Candida albicans* pellets (1:10 target cells), 400 µL of water-saturated phenol chloroform (ROTI®Aqua-P/C/I, Carl ROTH), and incubated for 1 hour at 65 °C at 1 400 r.p.m on a thermoshaker. Cell suspensions were cooled down on ice for 5 minutes. Samples were centrifuged at 13 000 g for 10 minutes at 4 °C, followed by transfer of the aqueous phase to an equal volume of water-saturated phenol and chloroform (ROTI®Aqua-P/C/I, Carl ROTH). For RNA precipitation, the aqueous phase was mixed with 500 mL cold isopropanol and 50 µL 3M sodium acetate (pH 5.2) and incubated at -20 °C overnight. The precipitated RNA was collected by centrifugation at 4 °C and washed twice with 75-80% ethanol, air-dried, and re-suspended in water. Isolated RNA was treated with DNase I (Qiagen) and further purified on column using RNeasy MinElute Cleanup Kit (Qiagen) according to the manufacturer’s instructions. Total RNA samples were submitted for Truseq Total RNA library preparation and sequenced by 100 bp single-read Illumina sequencing.

Reads were pre-processed to remove adapters and low-quality sequences and mapped to a reference consisting of Human (GRCh38.p13), *S. cerevisiae* (R64-1-1.98), and *C. albicans* (C_albicans_SC5314, Assembly22) genomes using STAR (version 2.7.10b) (https://github.com/alexdobin/STAR). Fraction of YAC-specific reads from total RNA reads was estimated excluding ribosomal RNA (rRNA) reads. Total read counts from *S. cerevisiae* were summarized and normalized to total *C. albicans* reads per sample. Concentration of transcripts per cell was performed by dividing the normalized *S. cerevisiae* read counts by the mean cell volume determined by Coulter counter measurements. Gene Set Enrichment Analysis (GSEA, version v4.3.2) and single-sample GSEA (ssGSEA) ^65^ were performed from *S. cerevisiae* aligned reads.

### Spike-in normalized Pol II ChIP-seq

To study the occupancy of RNA polymerase II on YAC-containing strains, we performed Spike-in normalized ChIP-seq ^39,40^. Yeast cultures from *Saccharomyces cerevisiae* and *Candida glabrata* (spike-in) were grown separately in YPD at 30 °C. *S. cerevisiae* cultures were mixed with *C. glabrata* at an OD_600_ ratio of 1:2 and immediately fixed (<5s) with the addition of 1% Formaldehyde (10-15% Methanol, Sigma-Aldrich) and incubated for 15 minutes at 30 °C. Fixed cultures were quenched with 0.125 M glycine for 5 minutes at 30°C, followed by two washes in ice-cold Phosphate-buffered saline (PBS). Cell pellets were collected by centrifugation and subsequently snap-frozen. Thawed pellets were lysed in FA lysis buffer (50 mM HEPES-KOH pH 8.0, 1 mM EDTA, 150 mM NaCl, deoxycholate, 1% Triton X-100, 1 mM PMSF, cOmplete Protease Inhibitor) using ceramic beads in a FastPrep 24 homogenizer (MP Biomedicals). Cell lysates were adjusted to 1 mL and sonicated twice for 8 minutes each (10 seconds on/20 s off) using a 1/8″ microtip (Qsonica Q500) in a −20 °C ethanol bath. Debris was pelleted, and the supernatant was collected for input or ChIP. For each ChIP sample, 40 µl of Protein G Dynabeads (Invitrogen) were blocked in PBS containing 0.5% Bovine Serum Albumin (BSA) for 40 minutes at room temperature, pre-bound with 10 µg of antibody (anti-Rpb1, 8wG16) in PBS for 40 minutes at room temperature, and then washed twice with PBS. Beads were then incubated with 0.5 ml of supernatant at 4 °C overnight. Beads were washed twice in FA lysis buffer and three times in high-salt FA lysis buffer (50 mM HEPES-KOH pH 8.0, 1 mM EDTA, 500 mM NaCl, 0.1% sodium deoxycholate, 1% Triton X-100, 1 mM PMSF), then eluted in ChIP elution buffer (50 mM Tris-HCl pH 7.5, 1% SDS, 10 mM EDTA) at 65 °C for 20 minutes. Simultaneously, 15 µl of input was directly combined with 115 µl of ChIP elution buffer. Crosslinks were reversed at 65 °C for 5 h, followed by RNase A (37 °C, 1 h) and proteinase K (65 °C, 2 h) treatments. DNA was purified using the ChIP DNA Clean & Concentrator kit (Zymo Research) and sequenced by paired-end (2 × 150 bp) Illumina sequencing.

Reads from the ChIP and Input samples were mapped to customized reference genomes using bowtie2 ^64^ (version 2.5.2). Depending on the analyzed strain, alignments were performed against a combined reference index containing the budding yeast (*S. cerevisiae*, version R64-1-1/sacCer3) genome, the exogenous spike-in (*C. glabrata*) genome, and the specific YAC sequence (YAC-L or YAC-S), or against a control index lacking the YAC sequence. The maximum paired-end fragment length parameter was set to 1000 bp to accommodate longer fragments. Following alignment, the mapping results were converted to BAM format, coordinate-sorted, and indexed using SAMtools ^66^ (version 1.19).

To quantify Pol II occupancy across *S. cerevisiae* (Sc) and YAC genomes, sequencing reads were normalized as follows:

Global Pol II Occupancy (**Fig. S4a**): To determine the binding efficiency per single DNA copy independently of YAC copy number, total ChIP read counts for Sc or YAC were normalized by their respective total Input read counts:

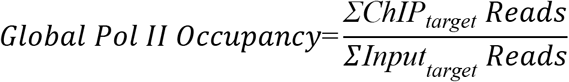

where target represents Sc or YAC reads.

Fraction of Pol II -bound YAC Reads (**Fig. 4d**): The fraction was calculated from the sum of the ChIP reads mapping to the YAC, normalized by the total ChIP reads:

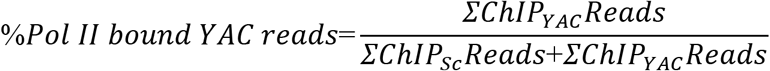

Normalized Pol II-bound Reads (**Fig. S4b**): To eliminate technical artifacts such as IP efficiency and sequencing depth variations, ChIP read counts were normalized by *C. glabrata* spike-in reads, adjusting for the initial chromatin mixing ratio derived from Input data:

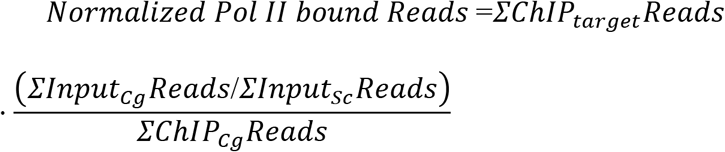

The same normalization factor was used to normalize ChIP-seq reads per genomic bin (**Fig. 4c, 4e).** Furthermore, to assess cellular Pol II-bound concentration (**Fig. 4f**), the sum of normalized Pol II-bound reads was divided by the respective cellular volume of each strain.

### SDS-PAGE and western blotting

For western blot analysis, cells were grown to exponential phase in YPD at 30 °C. The volume of 10 mL of cell culture was collected by centrifugation (2 minutes at 3,000 r.p.m.), resuspended in an equal volume of ice-cold 5% trichloroacetic acid (TCA, Roth), and incubated for 10 minutes on ice. Cell pellets were subsequently collected by centrifugation for 2 minutes at 3,000 r.p.m. at 4 °C and snap-frozen in liquid nitrogen. Collected samples were washed in 1 mL of acetone and air-dried. Subsequently, cells were lysed on a FastPrep-24 machine (MP Biomedicals) (6 cycles, 6.5 m/s, 45 s) in 100 µl lysis buffer (50 mM Tris-HCl pH 7.5, 1 mM EDTA pH 8.0, 50 mM dithiothreitol) containing 1x protease inhibitor cocktail cOmplete^TM^ (Roche) and using 100 µL volume of acid-washed glass beads (Sigma).

After lysis, samples were boiled in sample buffer (Tris-HCl pH 6.8, 30% Glycerol, 9% SDS, 6% β-mercaptoethanol, 0.05% Bromophenol blue) at 95°C for 5 minutes. Samples were loaded onto 7% Tris-acetate gels (Invitrogen) and resolved in 1x Tris-acetate SDS running buffer (Invitrogen). Proteins were transferred to PVDF membranes at 200 mA at 4°C for 2 hours using a wet transfer apparatus. For immunoblotting, membranes were blocked with 3% Milk in Tris-buffered saline with Tween 20 (TBS-T), before incubating with primary antibodies in 1% milk in TBS-T at 4°C overnight. Membranes were washed three times in TBS-T for 10 minutes each, incubated with secondary mouse IgG HRP-conjugated antibodies in 5% milk in TBST-T for 1 hour. Detection was performed using enhanced chemiluminescent substrate solutions SuperSignal West Pico and SuperSignal West Femto as described by the manufacturer (Thermo Fisher Scientific) and visualized on Fusion FX6 (Vilber). The primary and secondary antibodies used in this study are listed in Supplementary Table 5.

## Data availability

All sequencing data have been deposited in the European Nucleotide Archive (ENA) with the identifier PRJEB115124. All other raw data and reagents generated in this study are available upon request.

## End Notes

## Acknowledgements

We thank M. Peter for sharing reagents and laboratory infrastructure, S. Lüthi and S. Camenisch for laboratory support, S. Lee from ScopeM for assistance with microscopy experiments, F. Mair, R. Antonialli, I. Vgenopoulou, M. Malgorzata and A. Schütz from ETH Flow Cytometry Core Facility for technical assistance, S. Grüter from Functional Genomics Center Zürich (FGCZ) for analysis support, and A. Amodeo and Y. Barral for discussions and feedback. This work was supported by SNSF grants (PCEFP3_187003, 320030-232190) and an ETH Research Grant (ETH-38 21-1) awarded to G.E.N.

## Author Contributions

Conceptualization: M.E.E. and G.E.N.; Methodology: M.E.E., S.O., W.Q., and N.L.; Investigation: M.E.E., S.O., G.S., N.L., and S.S.L.; Writing – Original Draft: M.E.E. and G.E.N.; Writing – Reviewing & Editing: all authors; Funding Acquisition: G.E.N.; Supervision: G.E.N. and J.S.

---

**Supplementary Figure 1.**
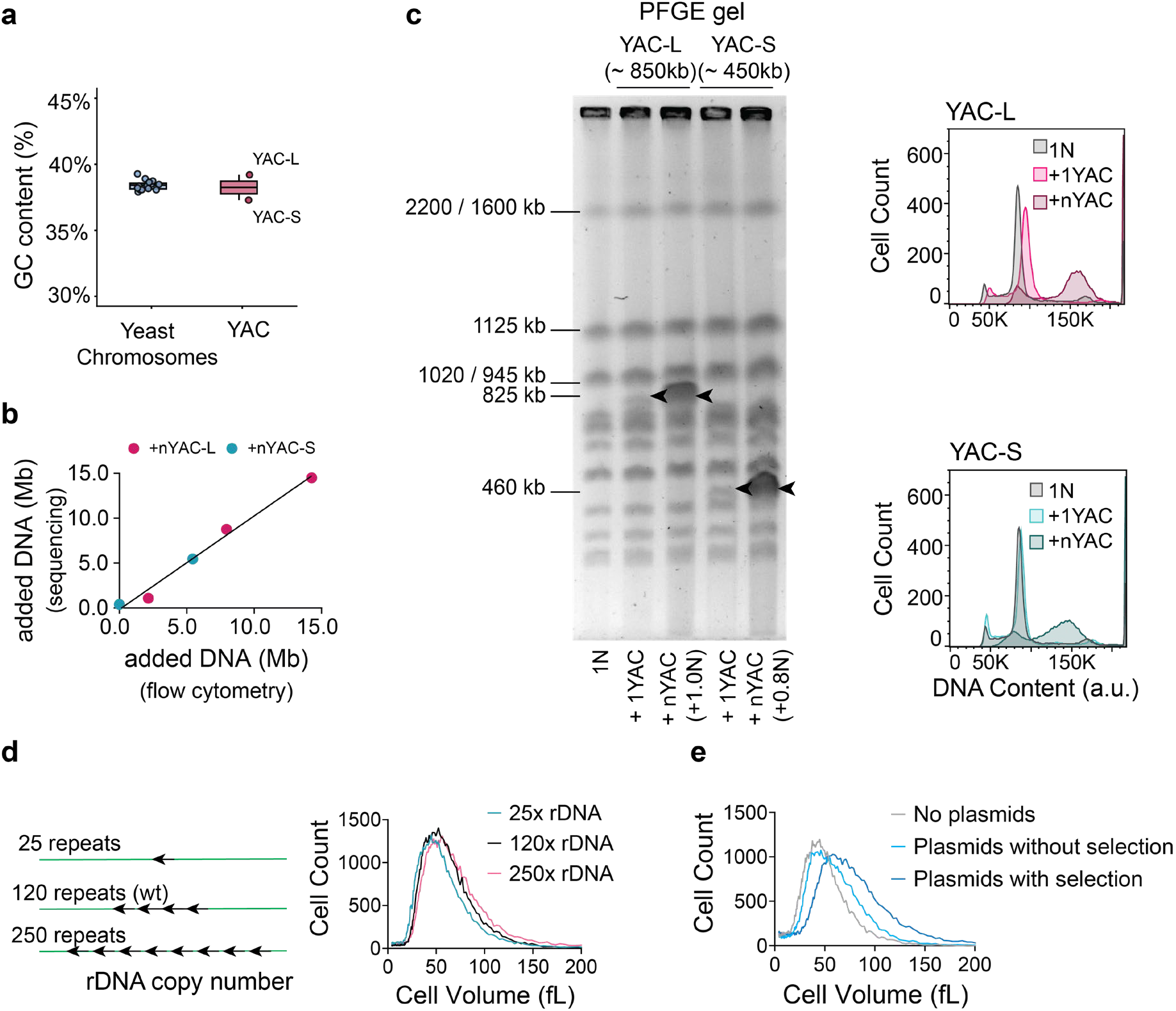
Experimental systems to increase non-coding DNA content. **a**, Nucleotide composition analysis (guanine-cytosine content) of *S. cerevisiae* genomic sequences and yeast artificial chromosomes YAC-L and YAC-S. **b**, Comparison between additional DNA estimated by nucleic acid staining assayed by flow cytometry and DNA sequencing from the experiments presented in Fig. 3a**-b**. DNA content by DNA sequencing was estimated normalizing YAC median read count to yeast chromosomes median read count of G2-phase samples. **c**, (*left*) Pulsed-field gel electrophoresis of isolated colonies with different amounts of accumulated YAC DNA. (*right*) DNA content analysis of these strains by flow cytometry. Representative of 2 biological replicates. **d**, The ribosomal DNA (rDNA) in budding yeast is located in a single locus that contains a variable number of the 9.1 kb tandemly repeated rDNA gene. Strains containing the indicated repeat copy number and a deletion of *FOB1* to stabilize rDNA repeat copy number were grown to exponential phase, and cell size was measured on a Coulter counter. For reference, 100 rDNA repeats correspond roughly to 10% of the yeast genome. Representative measurement of 3 independent experiments. **e**, Coulter counter volume measurements of cells carrying an 8.7 kb large 2µ multicopy plasmid (pGN442) containing a *LEU2* auxotrophic selection marker under a truncated promoter. This plasmid is present at an average of 40 copies per cell under non-selective conditions. Growth of these cells in medium lacking leucine (SC Leu-) selects for cells containing up to 200 plasmid copies ^67^, which corresponds to 20% of nuclear DNA.

**Supplementary Figure 2.**
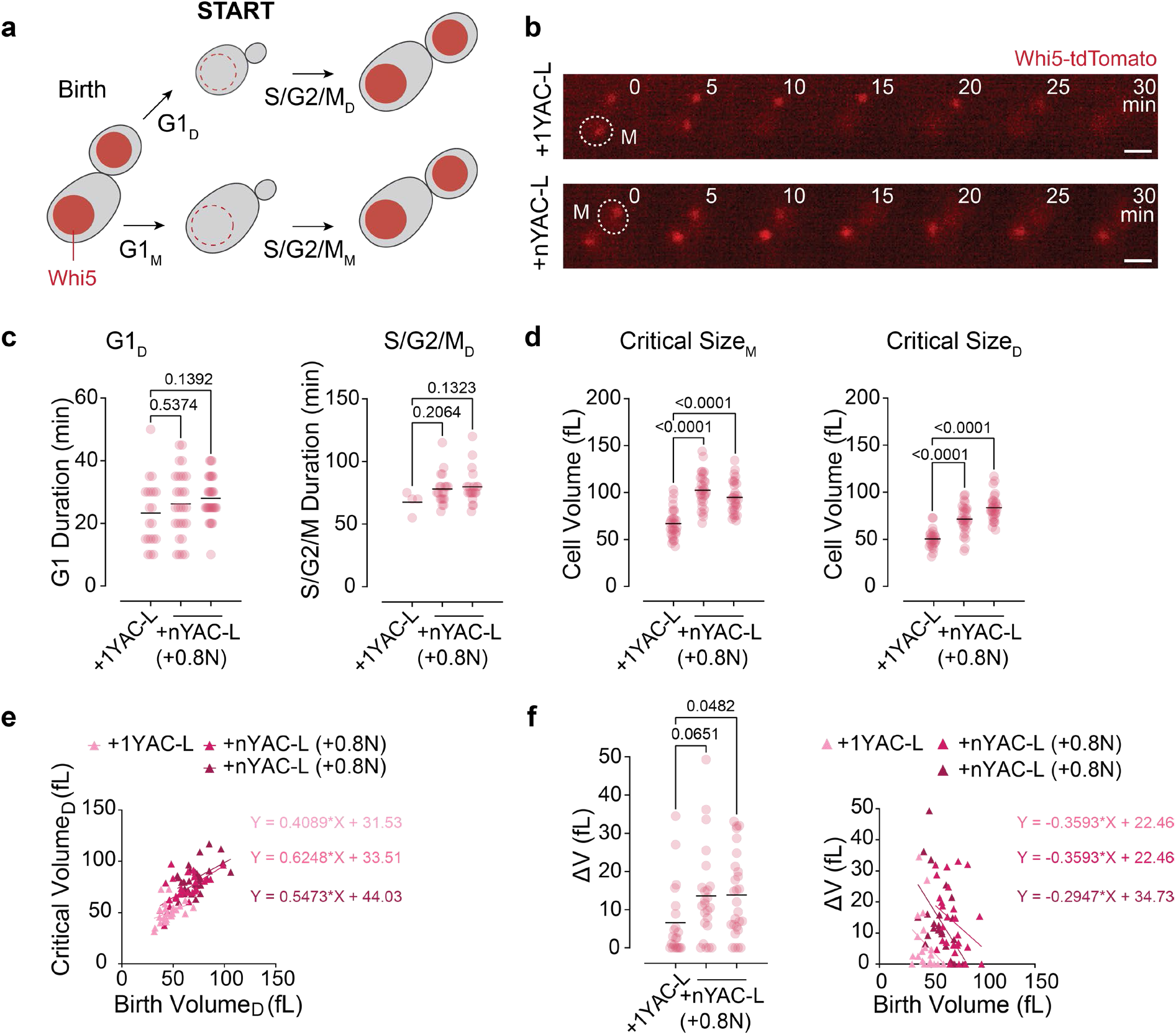
Non-coding DNA slows down cell cycle progression. **a**, Schematic of cell cycle progression analysis in cells expressing a Whi5-tdTomato fusion protein to determine the duration of G1-phase (Whi5 nuclear) and combined S/G2/M cell cycle phases (Whi5 cytoplasmic) in mother and daughter cells. **b**, Images of live cells expressing Whi5-tdTomato carrying different numbers of YACs. Scale bar 5 µm. **c-d**, Analysis of the same live-cell imaging experiment as shown in Fig. 2, carrying different numbers of YACs. **c**, G1 duration, and S/G2/M duration were measured in daughter cells (median, black line). Adjusted p values were calculated by Kruskal-Wallis test followed by Dunn’s multiple comparisons correction. **d**, the cell size at Whi5 export from the nucleus (critical size) was measured in mother and newborn daughter cells (mean, black line). >30 cells were analyzed per condition. Adjusted p values were calculated by one-way ANOVA followed by Dunnet’s multiple comparisons correction. **e**, Critical cell volume of strains carrying different numbers of YACs plotted against cell volume at birth. Linear regression was performed for the data of each strain separately. **f**, (*left*) Added cell volume during G1-phase of strains carrying different YAC copies (mean, black line). >30 cells were analyzed per condition. Adjusted p values were calculated by one-way ANOVA followed by Dunnet’s multiple comparisons correction. (*right*) Added cell volume during G1-phase of strains carrying different YAC copies plotted against cell volume at birth. Linear regression was performed for the data of each strain separately. The data from **c-f**, is generated from two independently isolated YAC-bearing strains. One representative dataset from two independent measurements is shown.

**Supplementary Figure 3.**
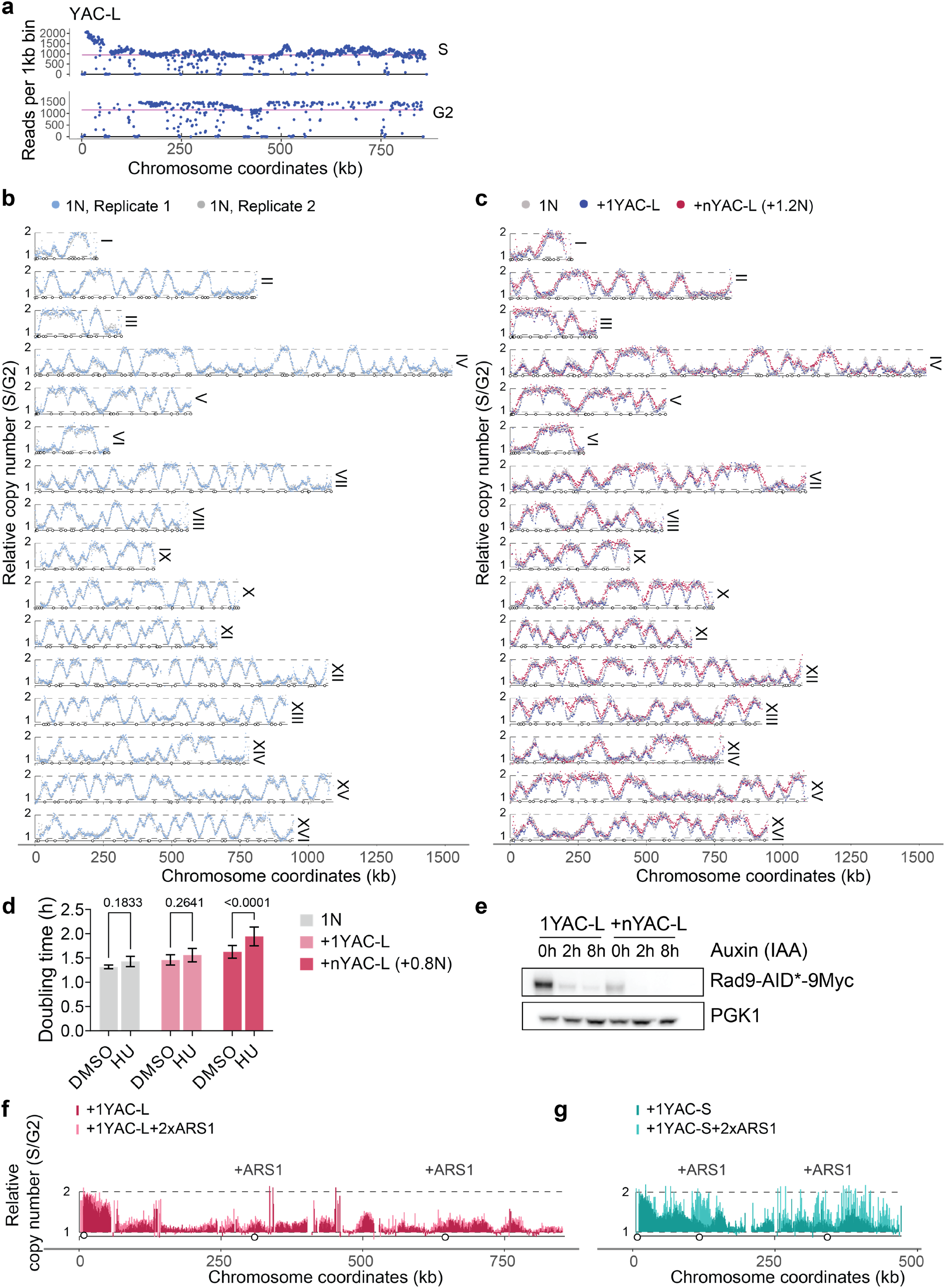
Replication analysis of DNA-burdened cells. **a**, YAC-L chromosome sequencing coverage of replicating (S) and non-replicating samples (G2). Data clusters close to the genome-wide median (pink line). **b**, Sort-seq replication profiles all *S. cerevisiae* chromosomes from haploid controls from two biological replicates. Circles indicate all annotated yeast origins of replication (ARS). Raw data were normalized to scale values between 1 and 2. The normalization factors (NF) are 1.456 for replicate 1 and 1.428 for replicate 2. **c**, Sort-seq replication profiles of *S. cerevisiae* chromosomes from haploid control (1N, gray, NF=1.456), a strain containing one YAC-L (1YAC-L: NF=1.415) or multiple YAC-L copies (nYAC-L: NF=1.494). Circles indicate annotated yeast origins of replication (ARS). **d**, Cell growth of the indicated strains was measured on a plate reader either in the presence or absence of 10 mM hydroxyurea (HU). Analysis of 3 independent experiments. Adjusted p values were calculated by two-way ANOVA followed by Šidák’s multiple comparisons test. **e**, The indicated strains expressing Rad9 fused with an auxin-inducible degron and a Myc-epitope tag were treated with auxin to induce Rad9 degradation (100 µM IAA). Samples were taken before and at 8 hours after auxin addition and subsequently processed for western blot analysis. PGK1 was used as a loading control. Representative of 2 biological replicates. **f**, **g**, Sort-seq replication profile of **d**, YAC-L with and without additional origins of replication sequence (*ARS1*) integrated at 300 and 630 kb (1YAC-L: NF=1.403; 1YAC-L:2x *ARS1*: NF=1.434), and **g,** YAC-S with added origins of replication (*ARS1* at 100 and 200kb) (1YAC-S, NF=1.451; 1YAC-S:2x ARS1, NF=1.387). Circles indicate annotated yeast origins of replication (ARS).

**Supplementary Figure 4.**
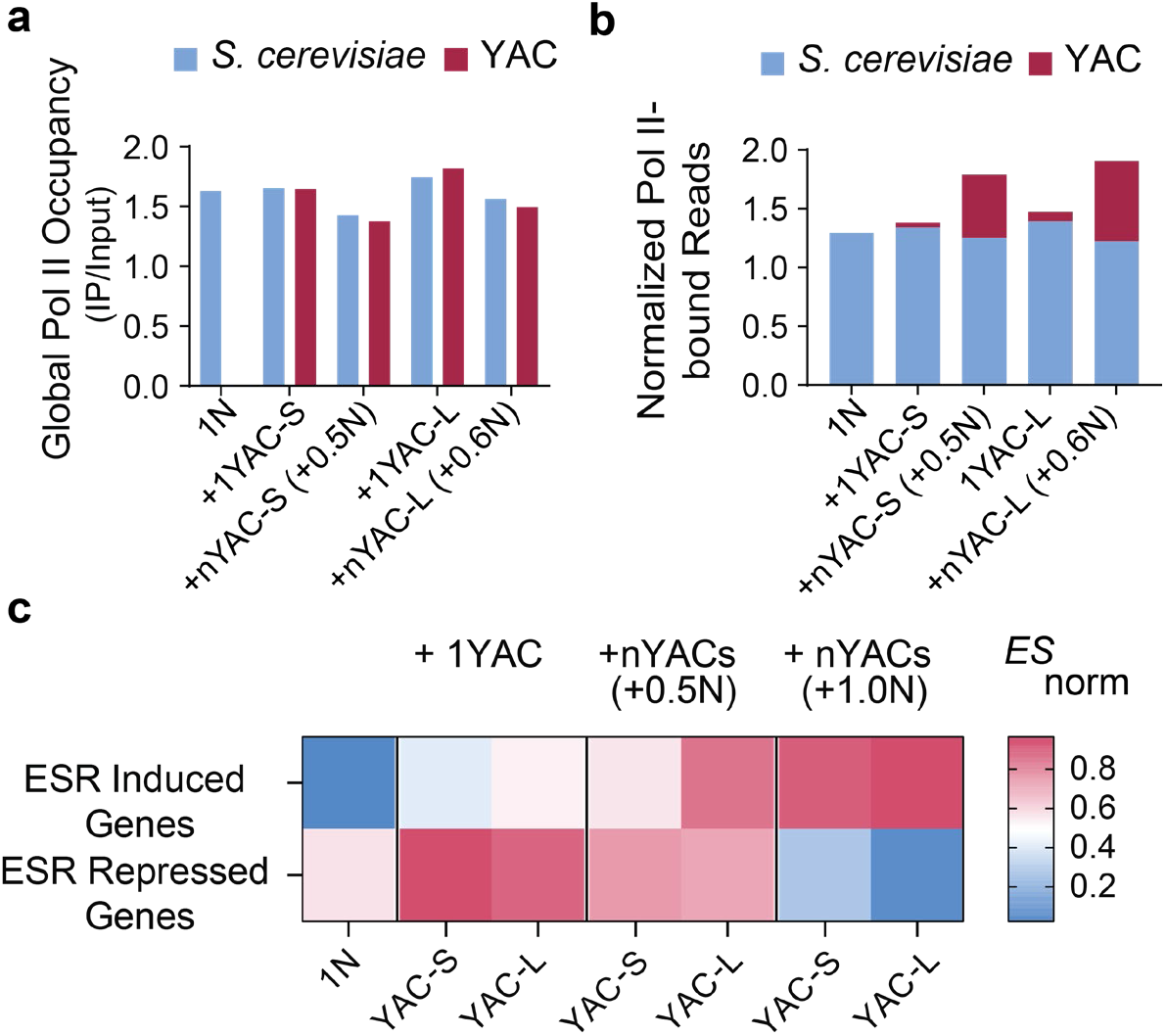
Pervasive transcription of YAC DNA impacts expression of yeast genes. a, Global Pol II occupancy per single DNA copy. Comparison of global Pol II binding efficiency per DNA copy on the *S. cerevisiae* and YAC sequences in the indicated strains. Values were determined by normalizing the sum of ChIP read counts by their respective sum of Input read counts (ChIP/Input). Signal from 2 pooled biological replicates per strain. See *Materials and Methods* for calculation details. **b,** Absolute quantification of Pol II binding. Stacked bar chart showing the absolute Pol II ChIP read counts mapped to the *S. cerevisiae* and YAC genomes. ChIP reads were normalized by *C. glabrata* spike-in and adjusted for initial chromatin mixing ratios derived from Input data. Signal from 2 pooled biological replicates per strain. See *Materials and Methods* for normalization details. **c,** Heat-map of single-sample gene set enrichment analysis (ssGSEA) of strains carrying different copies of YACs (YAC-L and YAC-S) for stereotypical environmental-stress response induced and repressed gene sets. Data for YAC-L and YAC-S bearing strains were combined from 2 biological replicates for this analysis.

**Supplementary Table 1.** *S. Cerevisiae* strain genotypes used in this study. These are all W303 background of genotype: ade2-1, leu2-3,112, ura3, trp1-1, his3-11,15, can1-100, GAL [psi+

| Number | Genotype |
| --- | --- |
| GN2089 | MATa, ade1::HIS3, lys2::KAN, pGAL1:CEN:NAT:YAC-2 (URA3/TRP1) |
| GN1407 | MATalpha, CLN2::CLN2p-GFPpest:HIS3, Whi5-tdTomato:KAN, pGAL1:CEN:NAT:YAC-2(URA3/TRP1) |
| GN902 | MATa, Whi5-tdTomato:KAN SPC42-GFP::TRP1 ADE2+ |
| GN2098 | MATalpha, Whi5-tdTomato:KAN, pGAL1:CEN:NAT:YAC-2(URA3/TRP1), SPC42-GFP::TRP1 |
| GN2103 | MATalpha, cln3::Hyg, pGAL1:CEN:NAT:YAC-2(URA3/TRP1) |
| GN1929 | MATa cln3::Hyg |
| GN2086 | MATalpha, lys2::KAN, pGAL1:CEN:NAT:YAC-2(URA3/TRP1) |
| GN2084 | MATalpha, ade1::HIS3, lys2::KAN, pGAL1:CEN:NAT:YAC-2(URA3/TRP1), cln3::mCitrine-CLN3-1-HIS3 |
| GN1682 | MATalpha, cln3::mCitrine-CLN3-1-HIS3 |
| GN2087 | MATa, pGAL1:CEN:NAT:YAC-2(URA3/TRP1) |
| GN2388 | MATa, YAC-7 (450kb), pGAL1:CEN:NAT:YAC-7 (URA3/TRP1) |
| GN3431 | MATa, YAC-7(450kb), pGAL1:CEN:NAT:terHIS3:ARS1(89kb):terKAN:ARS1(350kb):YAC-7(URA3/TRP1) |
| GN3440 | MATa, pGAL1:CEN:NAT:terHIS3:ARS1(300kb):terKAN:ARS1(630kb):YAC-2(URA3/TRP1) |
| GN1045 | MATalpha, CLN2::CLN2p-GFPpest:HIS3 Whi5-tdTomato:KAN fob1::HIS3 119xrDNA |
| GN1052 | MATalpha, CLN2::CLN2p-GFPpest:HIS3 Whi5-tdTomato:KAN fob1::KanMX ADE2+ 250xrDNA |
| GN1053 | MATalpha, CLN2::CLN2p-GFPpest:HIS3 Whi5-tdTomato:KAN fob1::HIS3 ADE2+ 27xrDNA |
| GN3276 | MATalpha, CLN2::CLN2p-GFPpest:HIS3, Whi5-tdTomato:KAN, leu2::pTEF1-osTIR::LEU2, RAD9-AID*-9Myc:HYG, pGAL1:CEN:NAT:YAC-2(URA3/TRP1) |
| GN3333 | MATa, Whi5-tdTomato:KAN, SPC42-GFP::TRP1, pGAL1:CEN:NAT:YAC-2(URA3/TRP1), leu2::pTEF1-osTIR::LEU2, RAD9-AID*-9Myc:HYG |
| GN1391 | MATa, ade1::HIS3, lys2::KAN, pGAL1:CEN:NAT:YAC-2(URA3/TRP1) |
| GN2450 | MATa, YAC-7 (450kb), pGAL1:CEN:NAT:YAC-7 (URA3/TRP1) |
| GN1391 | MATa, ade1::HIS3, lys2::KAN / pGAL1:CEN:NAT:YAC-2(URA3/TRP1) |
| GN2090 | MATa, ade1::HIS3, lys2::KAN /YAC-3, YAC-3 (670kb), pGAL1:CEN:NAT:YAC-3 (URA3/TRP1) |
| GN2094 | MATa, ade1::HIS3, lys2::KAN /YAC-7, YAC-7 (450kb), pGAL1:CEN:NAT:YAC-7 (URA3/TRP1) |

**Supplementary Table 2.** Original *S. Cerevisiae* strains containing YAC sequences.

| Number | YAC | DNA | Previous Nomenclature | Source |
| --- | --- | --- | --- | --- |
| GN1755 | YAC-L (850kb) | Human Chr. Y | yOX39, YAC-2 | Foote et.al., 1992 |
| GN1760 | YAC-S (450kb) | Human Chr. Y | yOX1, YAC-7 | Foote et.al., 1992 |

\*YACs have been re-sequenced for this study, and this sequencing data is available at the
European Nucleotide Archive (ENA) with the identifier PRJEB115124.

**Supplementary Table 3.** Plasmids used in this study.

| Number | Plasmid | Derived | Source |
| --- | --- | --- | --- |
| pGN442 | LexA-mCherry-pTOW | pTOW40836 | This study |
| pGN440 | pTOW40836 | pTOWug2-836 | Moriya et al., 2006 |

**Supplementary Table 4.** Chemicals and reagents used in this study.

| Chemicals and reagents | Source | Identifier |
| --- | --- | --- |
| 5-Fluoroorotic acid (5-FOA) | Thermo Fisher Scientific | R0811 |
| alpha-factor | GeneScript | RP01002 |
| SYTOX™ Green Nucleic Acid stain | Invitrogen | S7020 |
| RNAse A | Roche (Merck) | 10109169001 |
| Proteinase K | Panreac AppliChem | A3830,0100 |
| Zymolyase | Amsbio | 120491-1 |
| GelRed<sup>®</sup> Nucleid acid gel stain | Biotium | 41003 |
| Concavalin A | Sigma | C2010 |
| Thiolutin | Bioss | bs-82392c-5MG |
| Hydroxyurea | Sigma | H8627-10G |
| 3-Indoleacetic acid (IAA, auxin) | Sigma | I2886-25G |
| Phleomycin | Invivogen | ant-ph-2p |

**Supplementary Table 5.** Antibodies used in this study.

| Antibody | Source | Identifier |
| --- | --- | --- |
| Anti-Rad53 [EL7.E1] | abcam | ab166859 |
| Anti-MYC [9E10] | abcam | ab32 |
| Anti-PGK1 (22C5D8) | abcam | ab113687 |
| Anti-Rpb1 (8WG16) | Santa Cruz | sc-56767 |
| Anti-mouse IgG HRP | BioRad | 170-6516 |
| Anti-mouse IgG HRP | Merck | A6782 |

